# Environmental controls and habitat connectivity of phototrophic microbial mats and bacterioplankton communities in an Antarctic freshwater system

**DOI:** 10.1101/2020.07.13.200626

**Authors:** J. Ramoneda, I. Hawes, A. Pascual-García, T.J. Mackey, D.Y. Sumner, A.D. Jungblut

## Abstract

Freshwater ecosystems are considered hotspots of biodiversity in Antarctic polar deserts. Anticipated warming is expected to change the hydrology of these systems due to increased meltwater and reduction of ice cover, with implications for environmental conditions and physical connectivity between habitats. Using 16S rRNA sequencing, we evaluated the structure of microbial mat and planktonic communities within a connected watershed in the McMurdo Wright Valley, Antarctica to determine the roles of connectivity and habitat conditions in controlling microbial assemblage composition. We examined benthic and planktonic samples from glacial Lake Brownworth, the perennially ice-covered Lake Vanda, and the Onyx River, which connects the two. In Lake Vanda, we found distinct microbial assemblages occupying sub-habitats at different lake depths, while the communities from Lake Brownworth and Onyx River were structurally similar between them. Despite the higher connectivity between bacterial communities in the shallow parts of the system, environmental filtering dominated over dispersal in driving bacterial community structure. Functional metagenomics predictions identified genes related to degradation of halogenated aromatic compounds in surface microbial mats exposed to changes in water regimes, which progressively disappeared with increasing depth. Shifting environmental conditions due to increasing connectivity, rather than dispersal, may become the dominant drivers of bacterial diversity and functioning in Antarctic freshwater ecosystems.

## Introduction

Environmental change poses a major threat to the maintenance of terrestrial aquatic ecosystems, particularly due to alterations in hydrology (1). This is particularly acute for the cryosphere, where liquid water is primarily derived from melting snow and ice, and where winter-summer ice dynamics play a major role in driving ecosystem function. In the Antarctic Peninsula, significant increases in temperature over the last 50 years have corresponded with increases in melting of ice and liquid water supply (2). In the McMurdo Dry Valleys, complex processes have resulted in increases in lowland glacier melting (3, 4), which in turn has led to water level rise in endorheic lakes occupying the valley floors (5, 6).

Freshwater systems in the polar deserts of continental Antarctica are particularly important hosts of inland biodiversity, the bulk of which is microbial (7). In these meltwater-derived systems, primary productivity and biomass generation relies on phototrophic microbial communities, since allochthonous loadings of organic carbon are negligible from surrounding soils. To understand the implications of changing hydrology for biodiversity and ecosystem function in these lakes, it is important to investigate which environmental factors drive the spatial distribution and diversity of these microbial communities across habitats and depths (7, 8).

While a large number of microbial taxa are common across polar regions (9, 10), on a local scale variation in the taxonomic identity of microorganisms has been ascribed to environmental selection. A number of studies have investigated how Antarctic freshwater microbial diversity is affected by environmental drivers, in particular its responses to pH, salinity, light and water availability across a range of aquatic habitats (11, 12, 13, 14, 15, 16, 17). While most of these studies found links between environmental factors and microbial diversity and functionality at local scales, a significant amount of variation remains unexplained (18). For example, a study comparing samples from a range of depths in three lakes in the McMurdo Dry Valleys found high levels of similarity in benthic microbial mat community composition, with only a very low influence of irradiance and conductivity in structuring communities (19). The study ascribed this to wide environmental tolerance and slow turnover of most cyanobacterial taxa. It is therefore conceivable that factors related to the dispersal of bacterial propagules between habitats also play a role in defining local bacterial assemblages.

The globally unusual suite of conditions that characterize Antarctic freshwater ecosystems influence the degree of connectivity between habitats (20, 21). Water flow is strongly seasonal, and the endorheic (closed basin), meromictic lakes (perennially stratified lakes) have water columns stabilised by salinity gradients, within which there may be little, or unidirectional, connectivity (22). The Brownworth Lake – Onyx River-Lake Vanda (BOV) system in the McMurdo Wright Valley represents a good natural laboratory to evaluate the roles of connectivity and habitat in microbial community assembly. At its eastern end it contains a closed freshwater system formed by Lake Brownworth (an exorheic, proglacial lake), which is fed by seasonal runoff from the Lower Wright Glacier, the melt rate of which has increased in recent years (4). Lake Brownworth overflows seasonally into the Onyx River, which flows inland to discharge into the endorheic Lake Vanda, which is stratified by a salinity gradient (6; Fig. 1A, B). The BOV system thus provides a range of aquatic habitats, all of which are variously connected. The system is also subject to ongoing change, with flows in the Onyx River currently exceeding ablation from the lake surface, resulting in water level rise and the formation of new aquatic habitat at the lake margins. Colonisation of this new habitat further allows the importance of connectivity and filtering to be addressed.

**Figure 1:**
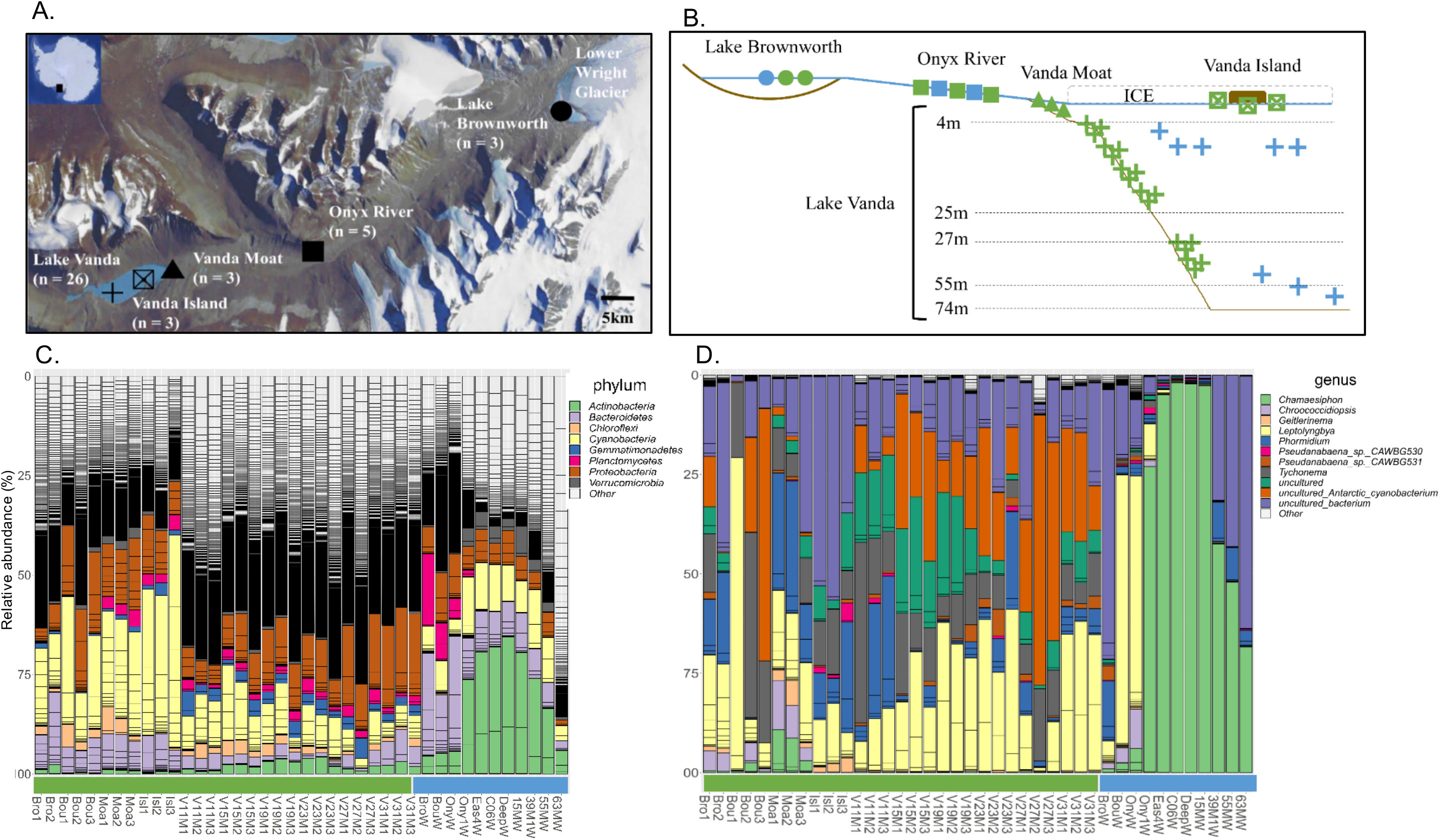
Location of the geographical units and bacterial community composition in the Wright Valley (Antarctica). (A) Aerial picture of the Wright Valley, from right to left: Lake Brownworth (circle), Onyx River (square), Vanda moat (triangle), Vanda island (open cross-square), and Lake Vanda (cross). (B) Cross-section of the sampling sites including the depth profile of Lake Vanda, divided by meromictic water layers. Green symbols = mats, blue symbols = water samples. (C) Relative abundance of dominant bacterial taxa (Phylum level), and (D) cyanobacterial taxa (Genus level), assessed using 16S rDNA sequencing. Samples are sorted left to right according to habitat (mat, in green; water, in blue) and by increasing depth.

In this study, we evaluated the bacterial diversity and community structure of the BOV system by sequencing the V4 region of the 16S rDNA gene of microbial mat and planktonic bacterial communities across geographical units and water depths. We hypothesized that benthic habitats with microbial mats, and planktonic habitats with bacterial communities in the water column would be compositionally distinct, but that local abiotic factors would strongly and consistently filter bacterial diversity. We also expected that the degree of connectivity between geographical units would correspond to similarities in bacterial community composition, and therefore the potential role of dispersal in structuring local assemblages was explicitly tested. By comparing benthic and planktonic communities, and covering shallow and deep environments, we address the role of dispersal and environmental filtering in structuring the bacterial communities. This is relevant for predicting how climate-driven hydrological changes will impact bacterial diversity in Antarctic freshwater ecosystems.

## Materials and Methods

### Study site and sample collection

The Wright Valley is located in the McMurdo Dry Valleys, in Southern Victoria Land, Antarctica (Fig. 1A, Fig. S1). Characterized by low annual average temperature (−19.8°C) and precipitation (below 100 mm year^−1^ water equivalent) (3), it is a closed freshwater basin. The low point of the valley is inland; glacial meltwater and groundwater flow towards this low point and evaporation and sublimation are the only water outputs from the system (23). Meltwater from the western side of the Lower Wright Glacier (77°25′S 163°0′E) feeds Lake Brownworth (77°26’4.96”S, 162°45’48.22”E, Fig. S1A and B), which is connected to Lake Vanda (77°31′S 161°34′E, Fig. S1C and F) through the seasonal Onyx River (Fig. S1D and E), a 32 km stream flowing through a polar desert landscape. Lake Vanda is perennially covered by a 3.5-4 m ice sheet, and in 2010 it was over 74 m deep (Hawes *et al*, 2013).

The structure of Lake Vanda reflects its current and historical climatic context, and is fully described in Castendyk et al. (3). In brief, the lake is understood to have undergone a series of climate-related filling and drying events (Fig. S2). The most recent high stand, some 50 m above the current level, was around 3000 years ago during a period of high meltwater production. Between 3000 and 2000 years ago, meltwater inflow declined and the lake evaporated to a shallow, hypersaline pool. Approximately 1000 years ago, renewed flow into the lake resulted in a freshwater layer from 27 to 55 m depth, mixed by double diffusion convection, overlying a salinity gradient. Additionally, approximately 100 years ago a new, discreet freshwater layer formed at the lake surface, which has since increased in thickness and now occupies the zone from 4 to 24 m depth. The lake continues to rise at approximately 0.25 m y^−1^ (3), and the shallowest depths sampled represent an inundation time series that can be dated from lake level records. The current lake structure is illustrated in Fig. S2.

A total of 29 cyanobacterial mat and 11 water samples were collected across the three main geographical units of the Wright Valley, namely Lake Brownworth, the Onyx River and Lake Vanda (collectively the BOV system, Fig. 1A and 1B, Table 1). The Lake Brownworth samples were collected in January 2012, and the remaining samples in December 2013. In addition to these three main geographical units, we categorized two additional shallow water subunits in which mat samples were collected also in December 2013 (Fig. 1B): The Vanda Moat, located at the interface between the Onyx River and Lake Vanda, and the Vanda Island.

**Table 1:** Sample types and water depths sampled at Lake Brownworth, Onxy River and Lake Vanda (McMurdo Wright Valley, Antarctica).

| Geographic location | Sample | Sample type | Water depth (m) | Replicates |
| --- | --- | --- | --- | --- |
|  |  | (Habitat) |  |  |
| **Lake Brownworth** | BRO | Mat | 0.1 | 2 |
|  | BROW | Water | 0.1 | 1 |
| **Onyx River** | BOU | Mat | 0.1 | 3 |
|  | BOUW | Water | 0.1 | 1 |
|  | ONYW | Water | 0.1 | 1 |
| **Lake Vanda moat** | MOA | Mat | 0.1 | 3 |
| **Lake Vanda moat 2017** | VMOAT | Mat | 0.1-1.9 | 3-5 per depth<br>(26 samples) |
| **Lake Vanda Island** | ISL | Mat | 0.1 | 3 |
| **Lake Vanda** | V11M | Mat | 11 | 3 |
|  | V15M | Mat | 15 | 3 |
|  | V19M | Mat | 19 | 3 |
|  | V23M | Mat | 23 | 3 |
|  | V27M | Mat | 27 | 3 |
|  | V31M | Mat | 31 | 3 |
|  | ONY1W | Water | 8 | 1 |
|  | EAS4W | Water | 15 | 1 |
|  | C06W | Water | 15 | 1 |
|  | DEEPW | Water | 15 | 1 |
|  | 15MW | Water | 15 | 1 |
|  | 39MW | Water | 39 | 1 |
|  | 55MW | Water | 55 | 1 |
|  | 63MW | Water | 63 | 1 |

The upper cohesive layer of the moat and river mats was sampled from the shore with a sterile spatula, and the deeper mats by a SCUBA diver using a 38 mm-diameter coring device from sites between 0.1-31 m deep. Mat samples were transferred into sterile plastic containers and immediately frozen at −20°C after collection, shipped frozen and stored at −80°C at the Natural History Museum (NHM, London, UK) until further processing. Water samples were collected from the shore (shallow samples from all sites) or using a Kemmerer water sampler through a hole in the ice of Lake Vanda (8-63 m). Biomass was collected from water samples by filtering 1-2 L through sterile Sterivex-GP^®^ filter units (Merck Millipore, Darmstadt, Germany) with 0.22 μm pore size. Biomass was stored in lysis buffer at −20°C immediately after processing, shipped frozen and stored at −80°C at the NHM until further processing.

Additionally, in January 2017 mat samples were collected from four shoreline sites at the eastern end of Lake Vanda, with the aim of obtaining more depth resolution in cyanobacterial mat community structure. At each site, a depth transect was obtained, sampling 3 mat samples at depths of 0.1, 0.5, 0.8, 1.1, 1.5 and 1.9 m using a 25 mm diameter rod-mounted piston corer. These depths correspond to inundation dates of 2017, 2016, 2014, 2013, 2012, 2011, 2010 based on lake level data in Castendyk et al. (3) and the McMurdo LTER data repository (http://mcm.lternet.edu/). The corer was cleaned with 70% ethanol between samplings. Cores were placed in 15 ml Falcon tubes and flash frozen in liquid nitrogen. Thereafter they were stored at −20°C until analysed four months after collection.

### DNA extraction, Polymerase Chain Reaction (PCR) and Illumina sequencing

DNA was extracted from 0.3-0.5 g of mat material using a MoBio PowerBiofilm DNA Isolation kit (Carlsbad, CA) following the manufacture’s protocol, and quantified using NanoDrop ND8000 (Labtech International, UK). The V4 variable region of the 16S rDNA gene was PCR-amplified using 8.84 μl of PCR grade water, 5 μl of 5x GoTaq Flexi buffer, 2 μl of 25 μM magnesium chloride (MgCl_2_), 0.8 μl of 20 mg/ml Bovine Serum Albumin (BSA), 0.16 μl of 200 μM dNTPs (deoxynucleoside triphosphate) and 0.2 μl of 5 μg/μl GoTaq polymerase. The reaction mix was completed with 1 μl of the 515F forward primer (10 μM) and 1 μl of the 806R barcoded reverse primer (24, 25; Table S1) (10 μM). For the samples collected in 2017, the same procedures were followed, except that the amplification was made with primers 341F and 805R (24, 26). A volume of 1 μl of template DNA completed the 20 μl reaction. PCR conditions involved an initial denaturation of 2 minutes at 94°C, followed by 45 seconds at 94°C, 1 minutes at 50°C of annealing temperature and 90 seconds at 72°C during 35 cycles. A final elongation of 72°C for 10 minutes followed and the products were kept at 10°C until removal.

The PCR products were visualised using electrophoresis in a 1% agarose gel. The amplicons were purified following an AxyPrep Mag PCR clean-up magnetic protocol (Axygen, New York, NY). The purified PCR amplicons were quantified using a Qubit 2.0 fluorometer (Life Technologies, Glasgow, UK). The multiplexed pooled amplicons were sequenced in an Illumina MiSeq platform at the Natural History Museum (NHM) sequencing facility.

### 16S rDNA gene sequence processing

The results from moat samples collected in January 2017 were processed separately from those taken in 2013. The majority of the sequence processing was conducted in QIIME (24, 25). For the 16S rDNA gene sequences, FastQC was used to verify the expected quality and read length of the initially demultiplexed 16S rRNA sequences. Primers and Illumina adaptors were trimmed using Cutadapt v1.14. The paired ends were stitched together using Flash v1.2.11, setting a minimum and maximum read overlap of 15 bp and 300 bp respectively, and a maximum mismatch density of 0.15. Sequences were quality and size filtered using Prinseq v0.20.4, with a maximum phred quality threshold of 20 and a size range of 250-300bp. The functions UPARSE and UTAX from USEARCH v9.2were used for OTU clustering, *de novo* chimera removal and taxonomic assignment using Silva (16S_128) as a reference database, setting a 97% similarity threshold and 80% sequence coverage. Chloroplast and mitochondrial sequences were excluded from the data using the *phyloseq* package v1.19.1 in R v3.5.2 (27). Sample coverage ranged between 96% and 99%, so diversity analyses were conducted on un-rarefied data (Fig. S3). A total of 4900431 16S rRNA sequences were finally processed, ranging from 60676 to 197711 sequences per sample, with an average (± SD) of 122511 (±25529) sequences. From the 2017 dataset, an average (± SD) of 7158 (±25528.84) sequences per sample were processed, totalling 372277 sequences. Before analysis of the diversity and community structure of these datasets, singletons were removed and only for beta diversity analyses sequence reads were relative abundance-transformed. The sequences were submitted to Genbank SRA (Bioproject ID: PRJNA638378).

### Analysis of the diversity of bacterial communities

The description of the composition, diversity and community structure of the BOV system bacterial communities was performed in R (27), with the packages *phyloseq* v1.19.1 (28), and *vegan* v2.5-5 (29)). We analysed the whole communities and also the diversity and composition of the cyanobacterial subset separately, given the latter’s importance for the productivity of the system. To estimate the alpha diversity, we considered the number of OTUs in each community (Observed species index), and the Chao1 and Simpson’s (D) diversity indices. Beta-diversity was estimated by computing the Bray-Curtis dissimilarity, and ordination was performed by computing Non-Metric Multidimensional Scaling (NMDS) and Constrained Correspondence Analysis (CCA). The ANOSIM test with 999 permutations was used to test the similarity between bacterial communities from different geographic units and habitats in the BOV system.

### Identification of environmental drivers of bacterial community structure

Relationships between measured environmental parameters (i.e. temperature, conductivity, and pH) and bacterial community structure were statistically assessed using an ANOVA-like permutation test on the CCA model, with 999 permutations. Using the *permutest.cca* function in *vegan v2.5-5* (29), the test computes the significance of the constraints on the ordination. In order to obtain mechanistic insights into the processes of assembly in the bacterial communities, we calculated the Mean Nearest Taxon Distance (MNTD) of each sample, and computed a z-score by comparing to a null distribution using the package *picante (v1.8.1)* (30). Briefly, the MNTD is the mean phylogenetic distance of each taxon to its closest relative in the community, and the comparison to a random draw of taxa from the same community gives a measure of phylogenetic dispersion. Z-scores higher than +2 indicate significant phylogenetic overdispersion and is interpreted as predominantly variable selection processes in driving community composition, while z-scores below −2 indicate significant underdispersion and predominance of homogenizing selection (31). Values falling between −2 and +2 (no phylogenetic signal) are indicative of ecological drift and dispersal. The input phylogenetic tree for the analysis was built from a multiple sequence alignment using MUSCLE (32).

### Prediction of bacterial metagenomes

We predicted the metagenomes from the 16S rRNA sequences using PiCRUST (33). The method uses reference genomes that are biased towards human-gut taxa, and hence predictions should be taken with caution in other environments. We addressed the quality of the prediction by computing the Nearest Sequenced Taxon Index (NSTI), which indicates the mean similarity of the relatives used in the prediction of a given community (e.g. a NSTI of 0.05 indicates a 95% similarity on average). In Langille et al. (33) the authors showed that the quality of the prediction decreases with increasing NSTI values depending on the environment. In particular, for soil environments (including 6 Antarctic samples taken from Fierer et al. (34), they showed that the accuracy of the prediction is high (Spearman correlation coefficients with respect to metagenome sequencing around 0.8) for NSTI values as high as 0.20. Indeed, previous predictions in mat samples were considered high quality for mean values of 0.11 (35). Our mean NSTI values were 0.18, which lie within the high-quality boundaries for mat samples, and we also verified that the NSTI values were uncorrelated to depth (Fig. S4). Nevertheless, special care was taken in the interpretation of water samples, especially for those in the deepest part of Lake Vanda exposed to higher salinity, which is known to decrease the accuracy of the prediction.

Predictions were first analysed by reducing the dimensionality using Principal Component Analysis (PCA) with STAMP v2.1.3 (36), to investigate which environmental variables were most closely associated with the clustering of the communities. We then merged the genes into genetic pathways within the highest resolution level in the KEGG (Kyoto Encyclopaedia of Genes and Genomes) classification (37), and tested which pathways showed significant differences across the identified environmental variables. To address this question, we tested if the mean proportions of genes belonging to a given pathway were significantly different between communities belonging to two different conditions (e.g. water vs. mat), performing Games-Howell tests. We considered significant those tests with Bonferroni-corrected P-values below 0.05 and effect sizes larger than 0.4.

## Results

### 16S rRNA gene composition of microbial mats and water habitats

6347 different bacterial OTUs were found across all microbial mat and water habitats investigated in the BOV system, of which 170 OTUs belonged the phylum Cyanobacteria. Alpha diversity results are presented in Table 2. In microbial mats, the highest bacterial richness was found in Lake Vanda (average Observed Species index ± SE = 1782 ± 136 OTUs), followed by Lake Brownworth (1591 ± 14 OTUs), and Onyx river (1525 ± 313 OTUs) (Table 2), while the highest cyanobacterial richness occurred in Lake Brownworth (65 ± 3 OTUs). In the water habitat Onyx river hosted on average five times more bacterial OTUs than Lake Vanda’s water column (Onyx river: 3098 ± 106 OTUs; Lake Vanda: 626 ± 302 OTUs), while the sample from Lake Brownworth had richness values closer to Onyx river (2112 OTUs) (Table 2). The same pattern was observed for Cyanobacteria.

**Table 2:** Average (±SE) Chao1 diversity estimate and Simpson’s diversity index per sample for bacterial and cyanobacterial microbial mat and water communities in the BOV (Brownworth-Onyx-Vanda) system.

| Geographical unit | Microbial mat |  |  |  | Water |  |  |  |
| --- | --- | --- | --- | --- | --- | --- | --- | --- |
|  | Prokaryotes |  | Cyanobacteria |  | Prokaryotes |  | Cyanobacteria |  |
|  | Chao1 | D | Chao1 | D | Chao1 | D | Chao1 | D |
| **Lake Brownworth (n=3)** | 1839 (37) | 0.987<br>(0.004) | 73 (2) | 0.843<br>(0.060) | 2464 | 0.961 | 78 | 0.576 |
| **Onyx river (n=5)** | 1802<br>(324) | 0.950<br>(0.025) | 69 (11) | 0.551<br>(0.123) | 3444<br>(98) | 0.969<br>(0.009) | 95 (4) | 0.596<br>(0.085) |
| **Vanda moat (n=3)** | 1725 (80) | 0.978<br>(0.003) | 67 (8) | 0.848<br>(0.037) | - | - | - | - |
| **Vanda island (n=3)** | 1112<br>(141) | 0.945<br>(0.003) | 52 (5) | 0.722<br>(0.076) | - | - | - | - |
| **Lake Vanda (n=18)** | 2038<br>(139) | 0.981<br>(0.008) | 51 (7) | 0.807<br>(0.086) | 849<br>(378) | 0.935<br>(0.016) | 39 (24) | 0.284<br>(0.232) |

At the phylum level, there were strong compositional differences between the microbial mat and water habitats, particularly in Lake Vanda, as well as a difference by depth. Dominance of Cyanobacteria and Proteobacteria in microbial mats contrasted with high relative abundance of Actinobacteria in water (Fig. 1C). Within the mat habitat, cyanobacterial abundance dropped from an average 29.1 % in shallow water samples less than 1 m depth (i.e. Brownworth-Onyx and the shores of Lake Vanda), to 11.8 % in the samples from Lake Vanda below ice cover (11-31m deep). This change in relative abundance did not reflect a clear compositional change at the cyanobacterial genus level, which was dominated by the filamentous oscillatorians *Phormidium, Tychonema* and *Leptolyngbya* (Fig. 1D). Within the water column, location and depth also played a significant role on bacterial taxonomic composition. At the phylum level, communities shifted from Bacteroidetes- and Planctomycetes-dominated assemblages in Lake Brownworth and Onyx river, towards Actinobacteria-dominated communities in samples under ice cover in Lake Vanda (Fig. 1C). A strong shift in cyanobacterial community composition was also observed, as water samples under ice cover were dominated by unicellular *Chamaesiphon*, while *Leptolyngbya* dominated shallow water samples (Fig. 1D).

### Comparison of bacterial community structure and environmental drivers

Microbial mat and water habitats contained very distinct bacterial assemblages (Fig. 2A). The community structure of microbial mat samples from shallow locations (i.e. Lake Brownworth, Onyx River, Vanda Island and Vanda Moat) clustered together, and all were more similar to their corresponding water samples than was the case in the deeper water column samples in Lake Vanda (Fig. 2A). The relationship between the measured abiotic factors (i.e. temperature, conductivity, and pH; see Table S1) and bacterial community structure was assessed using CCA. Habitat type (i.e. mat vs. water), and geographical location within the BOV system were the most important factors determining bacterial community structure (Fig. 2F). However, the constrained ordination, which explained 55.7% of variation in bacterial community structure in total (24.8% in the first 2 axes), identified temperature and conductivity as important factors of bacterial community variation, with no influence of water pH (Fig. 2B). Within Lake Vanda, the importance of depth, a covariate of the environmental factors, was shown by correlating NMDS1 and depth for microbial mat (Fig. 2C) and water (Fig. 2D) bacterial communities. This analysis revealed both mat and water communities gradually shift in structure with increasing depth. Finally, using the 16S rRNA gene analysis from the littoral zone of Lake Vanda, a sharp gradient in bacterial community structure was observed at a finer scale, within the first 2 m depth at Lake Vanda moat (Fig, 2E, and see Fig S5 for additional alpha and beta diversity results of the 2017 Vanda moat dataset).

**Figure 2:**
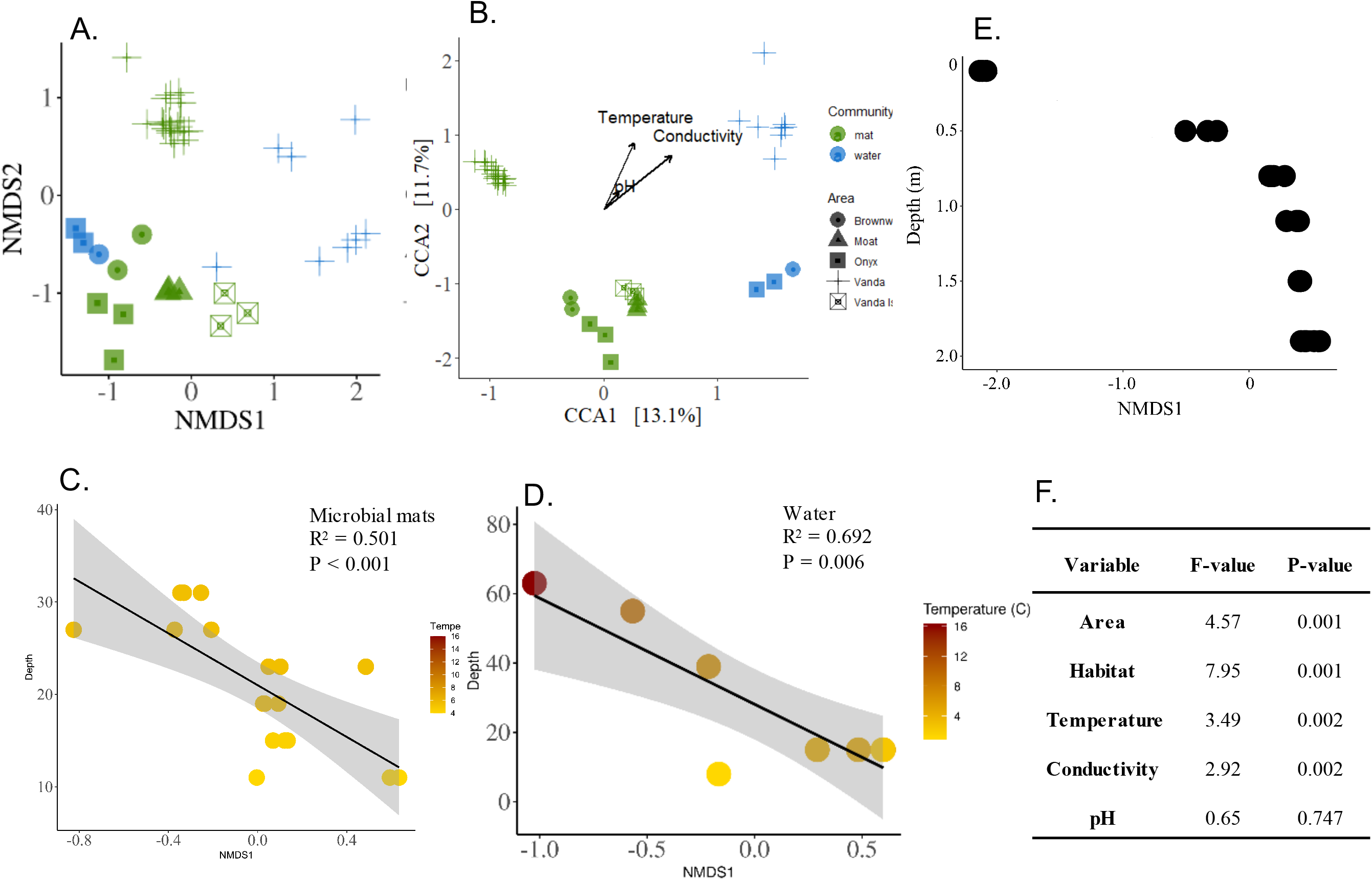
Beta-diversity patterns and bacterial and cyanobacterial communities in microbial mat and water habitats. (A) Non-metric multidimensional scaling (NMDS) ordination based on Bray-Curtis dissimilarities between bacterial communities across habitats and geographical units. (B) Constrained Correspondence Analysis (CCA) of bacterial community structure by temperature, conductivity and pH. Model outputs and statistical significance are reported in the table. Correlation between NMDS axis 1 (obtained from (A)) and depth within Lake Vanda, depicting the associated temperature gradient for microbial mat (C), and water (D) communities. (E) Correlation between NMDS axis 1 and depth within 2 m deep in Lake Vanda moat. The NMDS is reported in Figure S4.

### Potential role of dispersal between geographical units and habitats of the BOV system

In order to understand the potential role of dispersal between bacterial communities in the BOV system, a combination of the ANOSIM test and calculation of MNTD z-scores in different geographical units were performed. Microbial mat and water samples were classified into units by their physical connectivity (i.e. Lake Brownworth-Onyx river, Vanda Moat-Vanda Island, Lake Vanda at 8-15 m, Lake Vanda at 19-23 m, and Lake Vanda > 27 m) (Fig. 3C). The ANOSIM test revealed bacterial community structure was distinct at each of the habitat-geographical unit combinations (Fig. 3A), with the exception of the comparison between water communities of Lake Brownworth-Onyx river unit (n = 3), and Lake Vanda below 27 m deep (i.e. “Lake Vanda deep”, n = 3) (R = 1, P = 0.100), attributed to the small sample size of these sites. The MNTD randomization analysis supported the results of the ANOSIM, as the z-scores calculated in the different geographical units were equal to or lower than −2 (Fig. 3B), indicating a strong role for homogenising selection (31), and little or no influence of dispersal on the structure of the bacterial communities studied. Despite this clear observation, the value of the ANOSIM R was taken as an indicator of the potential dispersal of bacterial taxa between geographical units, and the potential dispersal pathways were shown in a diagram of the BOV system (Fig. 3C). This shows the most likely pathways of dispersal in the system, despite the influence of local environmental filters on the bacterial communities was more important than dispersal.

**Figure 3:**
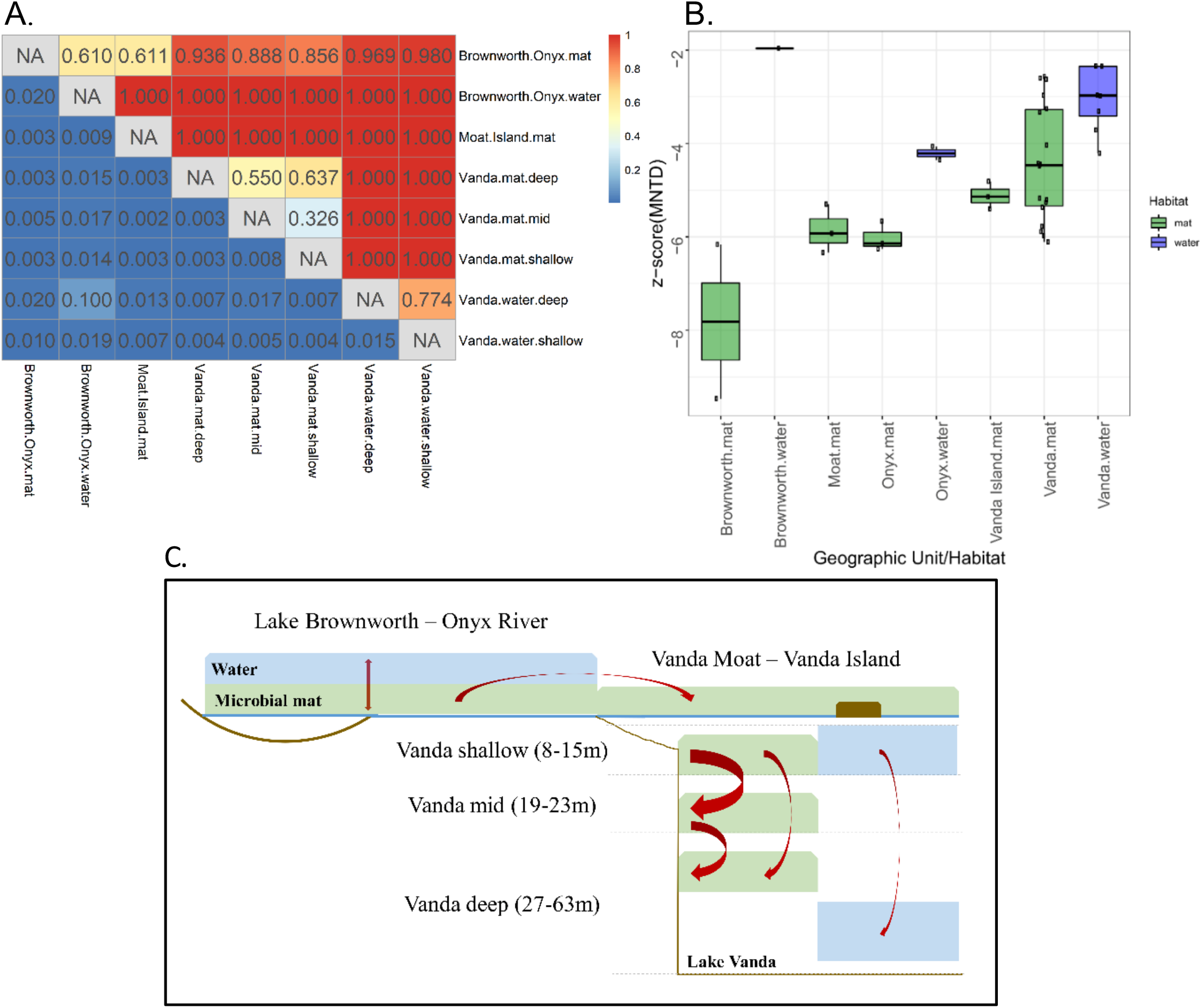
Test of bacterial community similarity and inference of environmental selection across habitats and geographical units in the BOV (Brownworth-Onyx-Vanda) system. (A) Analysis of similarities test (ANOSIM) between bacterial communities by geographical unit and habitat (mat vs. water) combinations. The upper half of the matrix depicts the ANOSIM R statistic, indicating higher community dissimilarity for values closer to 1; and the lower half depicts the P-values, with statistical significance set for P ≤ 0.05. (B) Mean Nearest Taxon Difference z-scores for bacterial communities across geographical units and habitats. Values below −2 are indicative of phylogenetic underdispersion within the community, indicating a role of homogenizing selection in driving community structure. (C) Schematic representation of the possible bacterial dispersal pathways between mat (green) and water (blue) habitats in the BOV system. Arrow thickness is inversely proportional to the R statistics reported in (A). R values above 0.8 were considered indicative of absence of dispersal.

### Metabolic function profiling

Predictions of metabolic pathways predominant in bacterial communities can point at potential bacterial adaptations to the environmental conditions, and reveal potentially unrecorded functions in natural communities. The gene inference analysis identified distinct metabolic functional profiles between microbial mat and water bacterial communities, which were strongly influenced by depth (Fig. 4A). The microbial mat and water environments were distinguishable by only a few metabolic functions (Fig. S6); instead, metabolic functional profiles for microbial mat communities gradually changed with increasing depth within Lake Vanda (Fig. 4B). The most important changes concerned functions associated to the degradation of halogenated aromatic hydrocarbon compounds, which characterized shallow mats (Fig. 4C-E). Deeper mat communities were characterized by predicted functions related to the synthesis of multiple amino acids and co-factors (Fig. S7).

**Figure 4:**
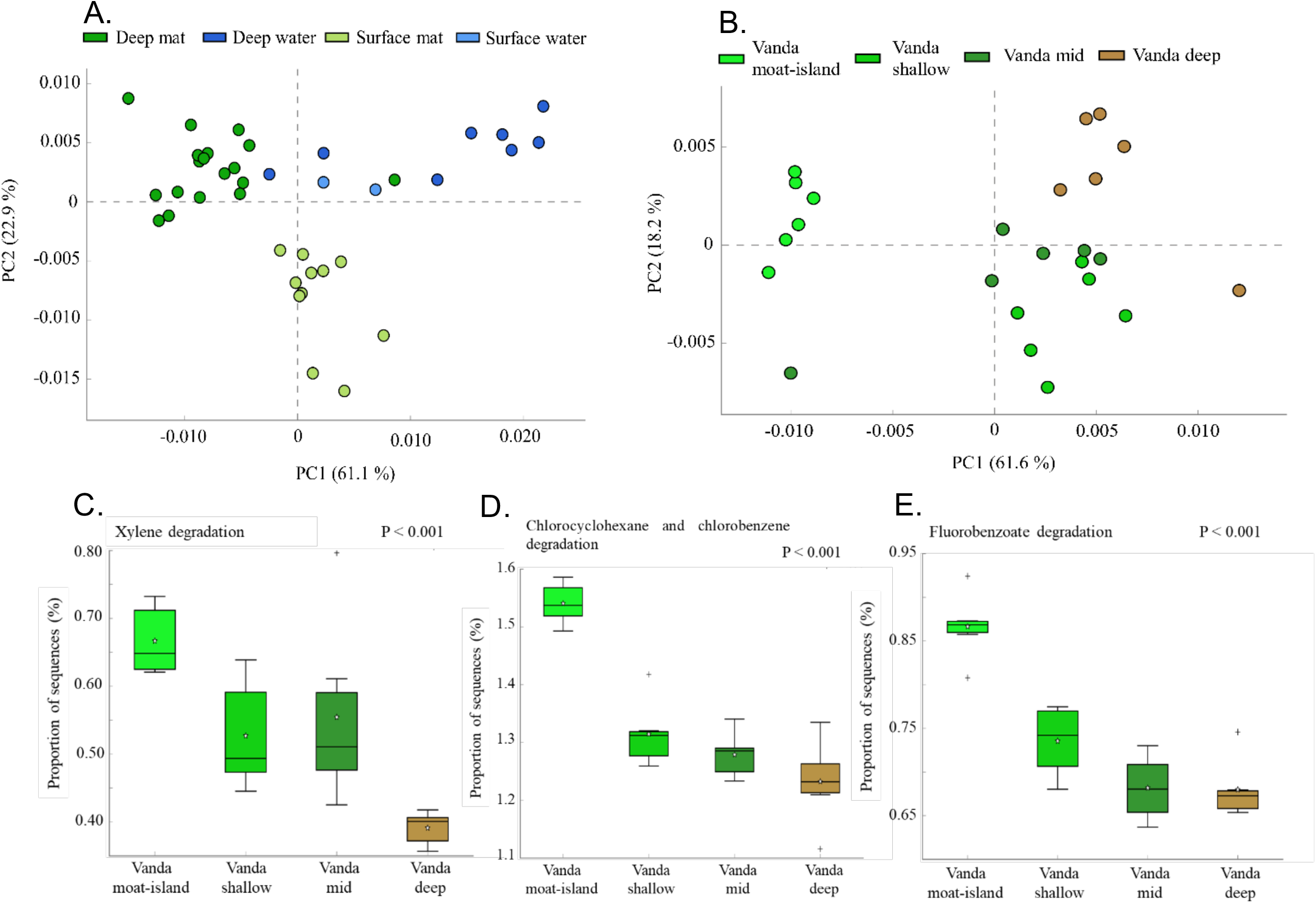
Principal component analysis (PCA) on the predicted bacterial metabolic gene composition between microbial mat and water communities at different depths in the BOV (Brownworth-Onyx-Vanda) system. (A) PCA on differences between the metabolic profiles of mat and water bacterial communities from the surface (<0.5 m) and deep (>4 m) environments, which correspond with different hydrological and temperature regimes. (B) PCA on differences between the metabolic profiles of bacterial communities in Lake Vanda, including its moat and island shores, and a division into three depth layers (shallow = 8-15 m, mid = 19-23 m, and deep = 27-63 m; for more information, see Fig. 3C). (C-E) Boxplots comparing the proportion of sequences predicting genes for xylene (C) and halogenated hydrocarbon compound degradation: (D) for chlorinated aromatic cycle degradation, and (E) for fluorobenzoate degradation.

## Discussion

In this study, we compared the 16S rRNA gene community structure and predicted metabolic functions of microbial mat and water bacterial assemblages across depths in an Antarctic freshwater system. Our results revealed a highly compartmentalized system in which high diversity is achieved through the strong environmental selection of taxa in different habitats.

Although some of these habitats are physically connected, bacterial dispersal between them has a limited role in structuring bacterial communities. This information is fundamental to address future increases in physical connectivity and changes in the environmental conditions in Antarctic freshwater systems due to ice meltdown (e.g. in Antarctic Lake Bonney, 38), which may promote strong shifts to the composition of microbial assemblages.

The composition and relative abundance of dominant bacterial and cyanobacterial taxa were very distinct between the microbial mat and water habitats in the BOV system (McMurdo Wright Valley, Antarctica). Dominance of Cyanobacteria, Proteobacteria and Bacteroidetes in the mats contrasted with a bacterioplankton dominated by Actinobacteria and with fewer Cyanobacteria. These observations agree with studies conducted in lakes Bonney Fryxell, Hoare and Miers (39, 40) suggesting differences between the bacterial benthos and plankton are widespread in Antarctic dry valley lakes. The cyanobacterial composition of the mats in the system was dominated at genus level by *Tychonema*, *Phormidium* and *Leptolyngbya*, which also agrees with previous studies on microbial mats in Southern Victoria land (19, 41).

Shallow parts of the system displayed higher structural similarity to each other than to sites under perennial ice cover, suggesting common responses to similar environmental conditions and a possible degree of dispersal among them. Indeed, the high (cyano)bacterial diversity detected in Onyx River suggests a central role of the river in propagule dispersal, also because Onyx river is the physical connector between lakes Brownworth and Vanda. The dominance of filamentous cyanobacterial *Leptolyngbya* in the water samples of Onyx river, most resembling those found in mat communities, could have originated from the mats in the Onyx river, and might have been suspended through disturbance by freeze-thawing and water currents during the rewetting of the dried mat in the river bed (42). Despite this role as the most likely dispersal pathway in the system, communities from Onyx river were markedly different from those under perennial ice cover in Lake Vanda, giving the view that despite bacterial propagule transport is commonplace in the shallow parts of the BOV system, propagules do not successfully establish in the locations they are transported to. The 2 m depth gradient observed for cyanobacterial mat communities in Lake Vanda moat further supports the idea that bacterial communities rapidly shift towards a common lake underwater assemblage. Considering the timespan each depth had been inundated, the convergence of sample composition at 1 m suggests that assemblages from the sampling locations on the shore of Lake Vanda achieved a common moat configuration within 3-4 years. Divergence of the most recently inundated communities from those below 1 m may reflect the early influence of terrestrial microbes pre-existing in soils, prior to colonists arriving from inflowing water and from deeper moat mats via mixing of moat water.

The possibility of dispersal as an effective driver of bacterial diversity in the BOV system was discarded by the finding that homogenising selection was the dominant driver of bacterial community structure, as has been reported in previous studies in vertically stratified lakes (11, 43). The observed gradual divergence of microbial mat and water communities with depth further supports the conjecture that deeper environments in Lake Vanda select for increasingly specialised bacterial assemblages (39). In the BOV system, an example of local habitat adaptation is the strong dominance of unicellular *Chamaesiphon* cyanobacteria, only recorded in the deeper layers of Lake Vanda. Dominant filamentous oscillatorian morphotypes had been recorded in Lake Vanda’s bottom of the euphotic zone (44), but not unicellular types. In Antarctica, *Chamaesiphon* had only been previously identified in microbial mats and sediment from glacial cryoconite holes in the Southern Victoria land and the McMurdo Dry Valleys (19, 45, 46). This novel observation highlights that very local selective processes determine the composition of bacterial assemblages in deeper layers of Antarctic lakes.

Strong environmental filtering should reflect on bacterial adaptations to the specific habitats under study. Prediction of metabolic functions with PiCRUST (33) gave insights into the different potential metabolic pathways across habitats and depths. The most important change in the predicted metabolic functionality for microbial mats happened with increasing lake depth in Lake Vanda. Pathways for the anaerobic degradation of xenobiotic halogenated aromatic hydrocarbon compounds characterized shallow mats, which is an important observation. While anaerobic oxidation of acetate for sulphur reduction is characteristic of the light and oxygen-depleted lower microbial mat layers (40, 47), these particular pathways had never been predicted in living Antarctic microbial mats. A recent metagenomic survey on microbial paleomats buried in soils around Lake Vanda identified proportions below 1% of xenobiotics metabolism genes (48). Since these functions are more commonly found in soil-borne bacteria (49), a plausible explanation is that shallow microbial mats may be composed of a number of soil-borne taxa recruited from the sediment and surrounding soil environment, which would explain the gradual decrease with depth. An alternative is that these functions are remnants of microbial activity happening after lake contamination by oil burning in the 1970s and 1990s (50). Future work should investigate whether alternative anaerobic metabolisms such as the ones predicted here are prevalent in Antarctic microbial mats, or whether these are indicative of past human impact.

## Conclusions

Environmental change is expected to lead to a redistribution of water and its availability in the McMurdo Dry Valleys (4), which will impact the connectivity between microbial habitats. This study shows that the Wright Valley contains distinct benthic and planktonic bacterial communities, which are primarily structured through environmental filtering. While superficial water flow seems to play a role in dispersing bacterial taxa, strong environmental filtering is a dominant process and leads to a vertical structuring of bacterial communities. Likewise, we show that within 3-4 years of inundation, bacterial communities achieve a common configuration in the shallow parts of Lake Vanda. These observations suggest that future impacts of hydrological changes on the bacterial communities will likely be driven firstly by changes in the physicochemical water properties, followed by an increasing role of connectivity as system stratification disappears. This implies that a homogenization of the water conditions could rapidly compromise the persistence of bacterial communities not adapted to those conditions. In Antarctica, understanding the effects of future environmental change on freshwater microbial diversity requires a detailed knowledge of how hydrological changes disrupt the diverse suite of environmental conditions that sustain bacterial diversity in the present.

## Acknowledgements

The authors acknowledge assistance by the sequencing facilities of the Natural History Museum London. Field research was funded by the New Zealand Ministry of Business, Innovation and Employment (grant UOWX1401 to IH). APG was funded by the Simons Collaboration: Principles of Microbial Ecosystems (PriME), award number 542381. The field work and logistics were supported by Antarctica New Zealand. We thank Hannah Christenson and Devin Castendyke for their help with the field work.

## Compliance with ethical standards

### Conflict of interest

The authors declare that they have no conflict of interest.

## Notes

### Competing Interest Statement

The authors have declared no competing interest.

