## Supplementary Material for "Environmental controls and habitat connectivity of phototrophic microbial mats and bacterioplankton communities in an Antarctic freshwater system"

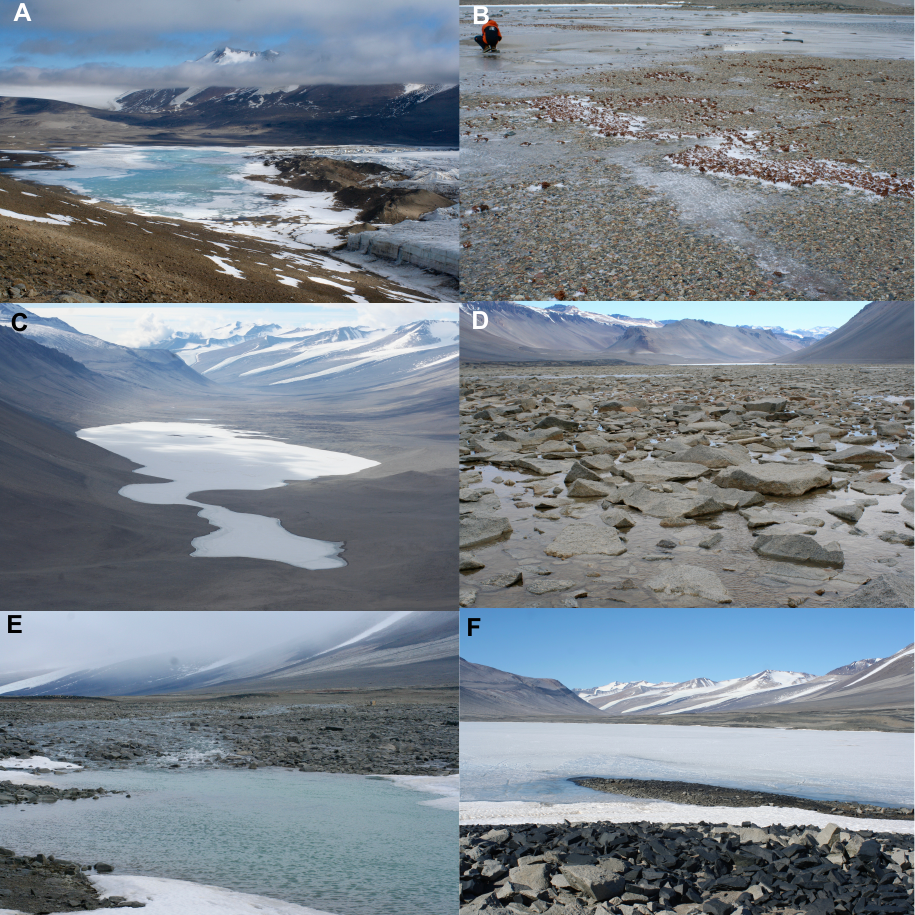

Supplementary Figure S1: Images showing A) Lake Brownworth, B) littoral zone of Lake Brownworth, C) Lake Vanda, D) Boulder Pavement at Onxy River, E) moat of Lake Vanda at the delta of Onyx River, and F) island in Lake Vanda with shallow littoral zone.

Onyx River

Lake Vanda

0 m

4 m

25 m

27 m

55 m

74 m

**Layer 1 - Moat**: Succession from soil-dominated to aquatic organisms, frequent freeze-drying cycles, expected similarity to the river.

**Layer 2**: ~100 years old. Constant chemistry and only light changes with depth. Any gradient will be due to light or age within the layer.

**Transition layer**: Expected to receive the taxa from above that are best adapted to moderately low light, low nutrients, and prolonged winter darkness.

**Layer 3**: ~1000 years old. Constant chemistry and warm (6°C). Only light changes with depth. Any gradient will be due to light or age within the layer.

**Layer 4**: ~1000-2000 years old. Linear increase in salinity with depth. Very warm (20°C). Considered the ancestral state of the lake ~2000 years ago.

Substrate

Water

ICE

Supplementary Figure S2. Detailed description of the depth profile of Lake Vanda, and the characteristics of its meromictic water layers.

| Sample | Location | GPS cords. | Sample type | Reps. | Depth (m) | Temp. (°C) | pH | | Conductivity (µS/cm) | |
| --- | --- | --- | --- | --- | --- | --- | --- | --- | --- | --- |
| BRO | Lake Brownworth | S77°26'4.96"  E162°45'48.22" | Mat | 2 | 0.1 | 1.0 | | 6.10 | | 38.7 |
| BOU | Onyx River’s boulder pavement | S77 31'22.4"; E161 45'34.1" | Mat | 3 | 0.1 | 1.7 | | 6.12 | | 57.0 |
| MOA | Onyx River moat | S77 31'24.2"; E161 41'17.3" | Mat | 3 | 0.1 | 1.6 | | 6.57 | | 77.1 |
| ISL | Lake Vanda central island | S77 31'36.6"; E161 36'56.9" | Mat | 3 | 0.1 | 1.3 | | 6.17 | | 159.7 |
| V11M | Lake Vanda | S77°32'15.48"  E161°32'42.34" | Mat | 3 | 11 | 4.0 | | 6.46 | | 970.0 |
| V15M | Lake Vanda | S77°32'13.27"  E161°32'5.74" | Mat | 3 | 15 | 4.4 | | 6.50 | | 985.0 |
| V19M | Lake Vanda | S77°32'12.71"  E161°32'41.69" | Mat | 3 | 19 | 5.5 | | 6.50 | | 1000.0 |
| V23M | Lake Vanda | S77°32'9.81"  E161°32'8.94" | Mat | 3 | 23 | 5.5 | | 6.50 | | 1100.0 |
| V27M | Lake Vanda | S77°32'10.35"  E161°33'17.65" | Mat | 3 | 27 | 5.5 | | 6.50 | | 1150.0 |
| V31M | Lake Vanda | S77°32'4.81"  E161°32'36.54" | Mat | 3 | 31 | 5.5 | | 6.50 | | 1260.0 |
| BROW | Lake Brownworth | 77°26'4.96"S, 162°45'48.22"E | Water | 1 | 0.1 | 1.0 | | 6.10 | | 38.7 |
| BOUW | Onyx River’s boulder pavement | S77 31'22.4"; E161 45'34.1" | Water | 1 | 0.1 | 1.6 | | 6.24 | | 55.6 |
| ONYW | Onyx River | S77 31'24.0"; E161 41'25.8" | Water | 1 | 0.1 | 1.2 | | 6.10 | | 78.5 |
| ONY1W | Onyx River moat | S77 31'28.9"; E161 40'53.2" | Water | 1 | 8 | 1.2 | | 6.23 | | 945.0 |
| EAS4W | Lake Vanda | S77 31'22.9"; E161 37'33.2" | Water | 1 | 15 | 4.8 | | 7.02 | | 993.0 |
| C06W | Lake Vanda | S77 31'38.9"; E161 36'16.0" | Water | 1 | 15 | 4.6 | | 7.27 | | 1006.0 |
| DEEPW | Lake Vanda | S77 31'59.6"; E161 32'02.2" | Water | 1 | 15 | 2.4 | | 7.11 | | 1033.0 |
| 15MW | Lake Vanda | S77°32'18.26"; E161°31'27.21" | Water | 1 | 15 | 4.4 | | 6.46 | | 985.0 |
| 39MW | Lake Vanda | S77°32'18.26"; E161°31'27.21" | Water | 2 | 39 | 6.5 | | 5.57 | | 1465.0 |
| 55MW | Lake Vanda | S77°32'18.26"; E161°31'27.21" | Water | 1 | 55 | 9.0 | | 7.48 | | 4280.0 |

Supplementary Table S1. Water physochemical properties of the samples from the different geographical units of the Wright Valley (Antarctica).

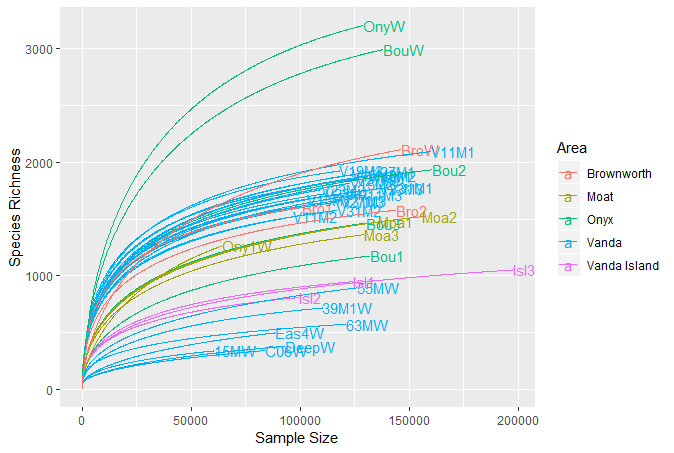

Supplementary Figure S3. Rarefaction curves of the samples from the different geographical units of the Wright Valley (Antarctica).

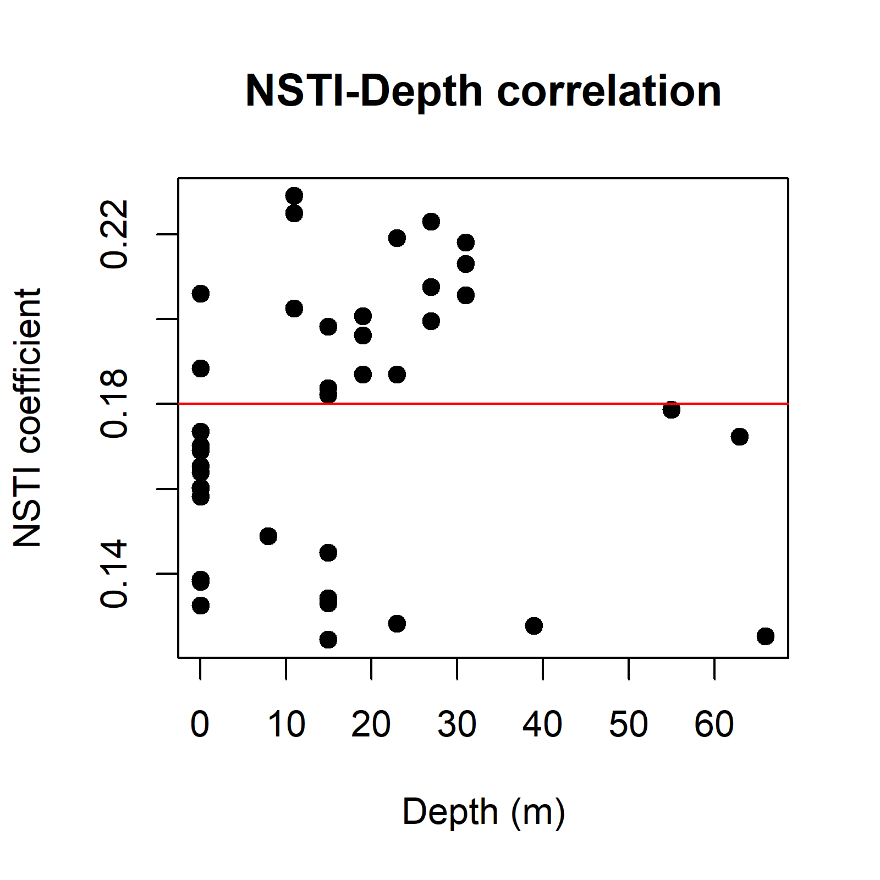

Supplementary Figure S4. Correlation between the Nearest Sequenced Taxon Index (NSTI) and depth of samples from mat and water habitats in the Wright Valley. The plot shows that there is no significant correlation between the two factors (r = 0.075, P = 0.646), implying no clear bias was observed among samples of deep and surface environments. The horizontal red line depicts the average NSTI value.

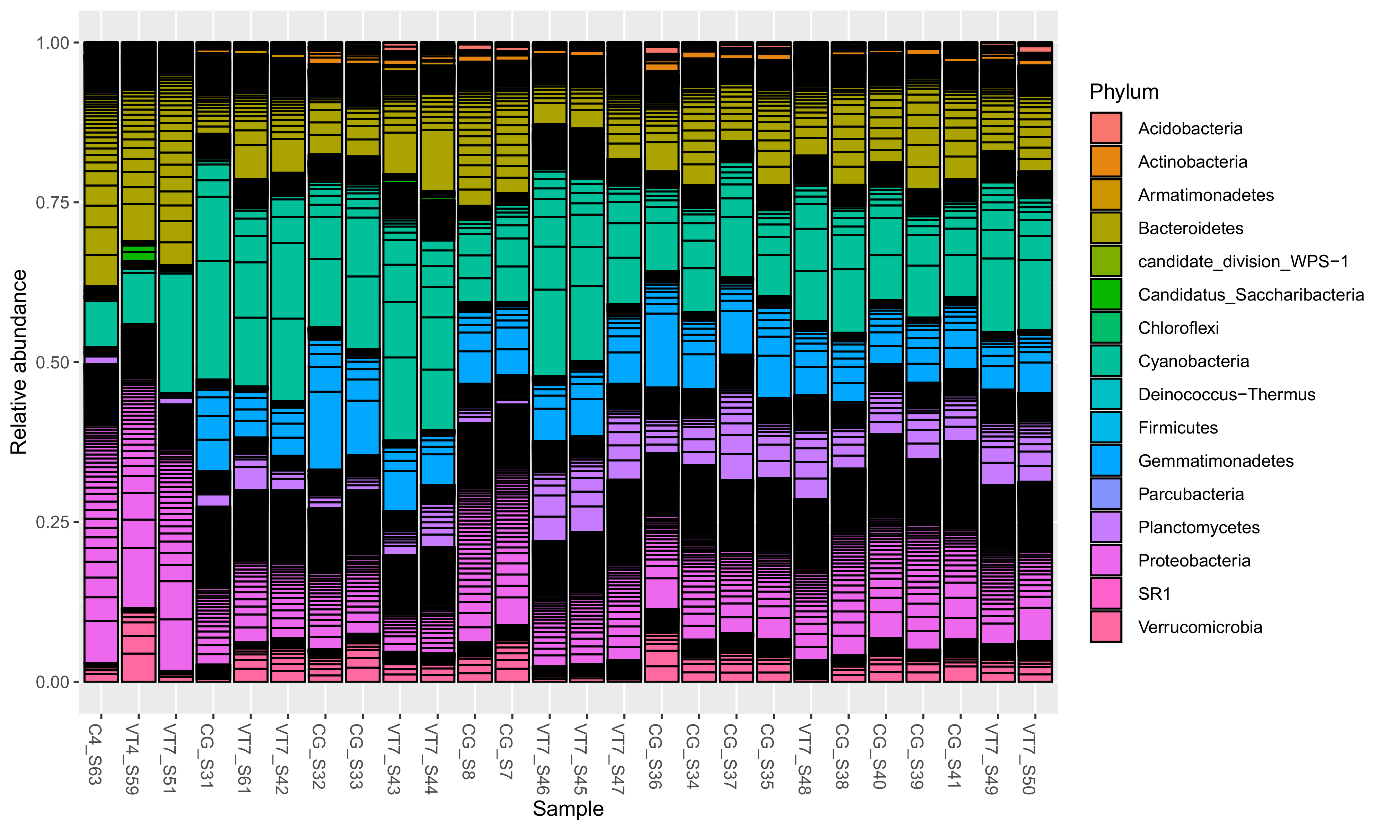

Supplementary Figure S5. Bacterial community composition, alpha diversity (Chao1 and Simpson’s diversity indices) and Non-Metric Multidimensional Scaling (NMDS) of the microbial mat communities of Lake Vanda moat within the first 1.9m deep at sites close to the entry of the Onyx River. Bacterial communities were described by sequencing the V3-V4 region of the 16S rRNA gene and clustering of OTUs at 97% sequence similarity.

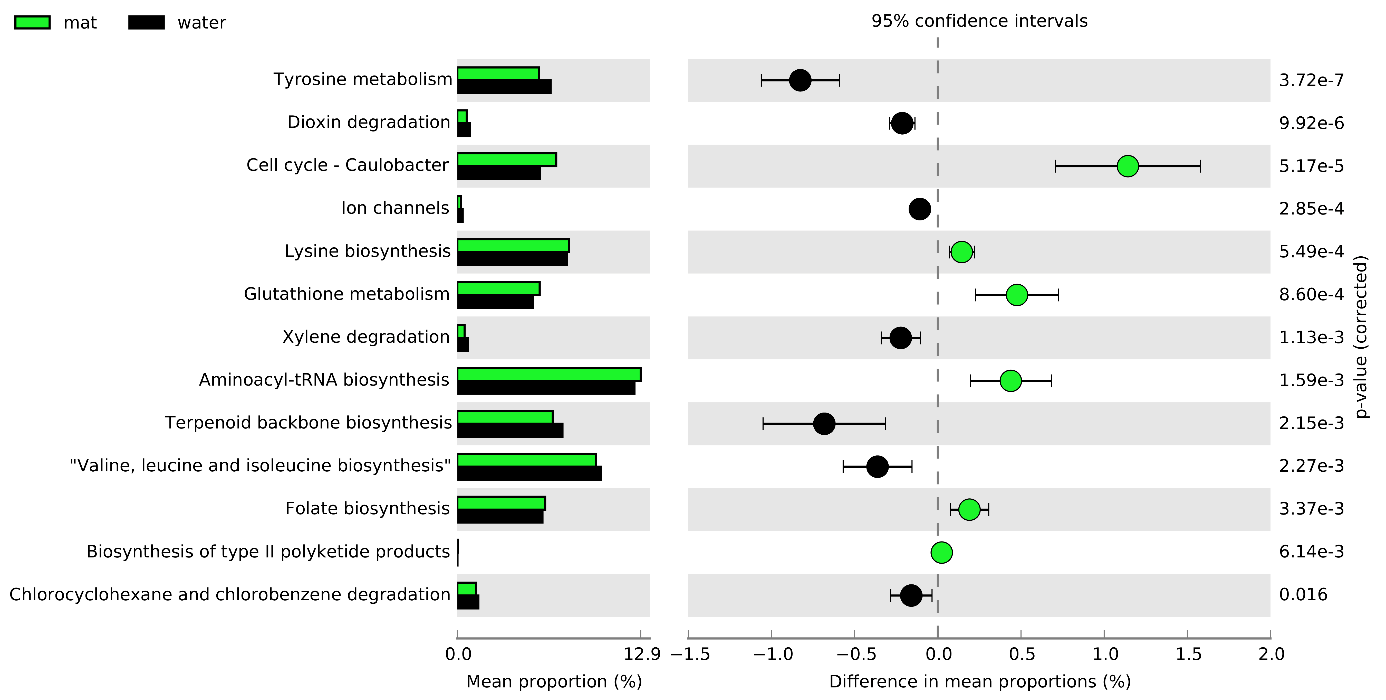

Supplementary Figure S6. Complete list of differences in the metabolic profiles comparing inferences for mat versus water bacterial communities. Only statistically significant differences are displayed (P < 0.05).

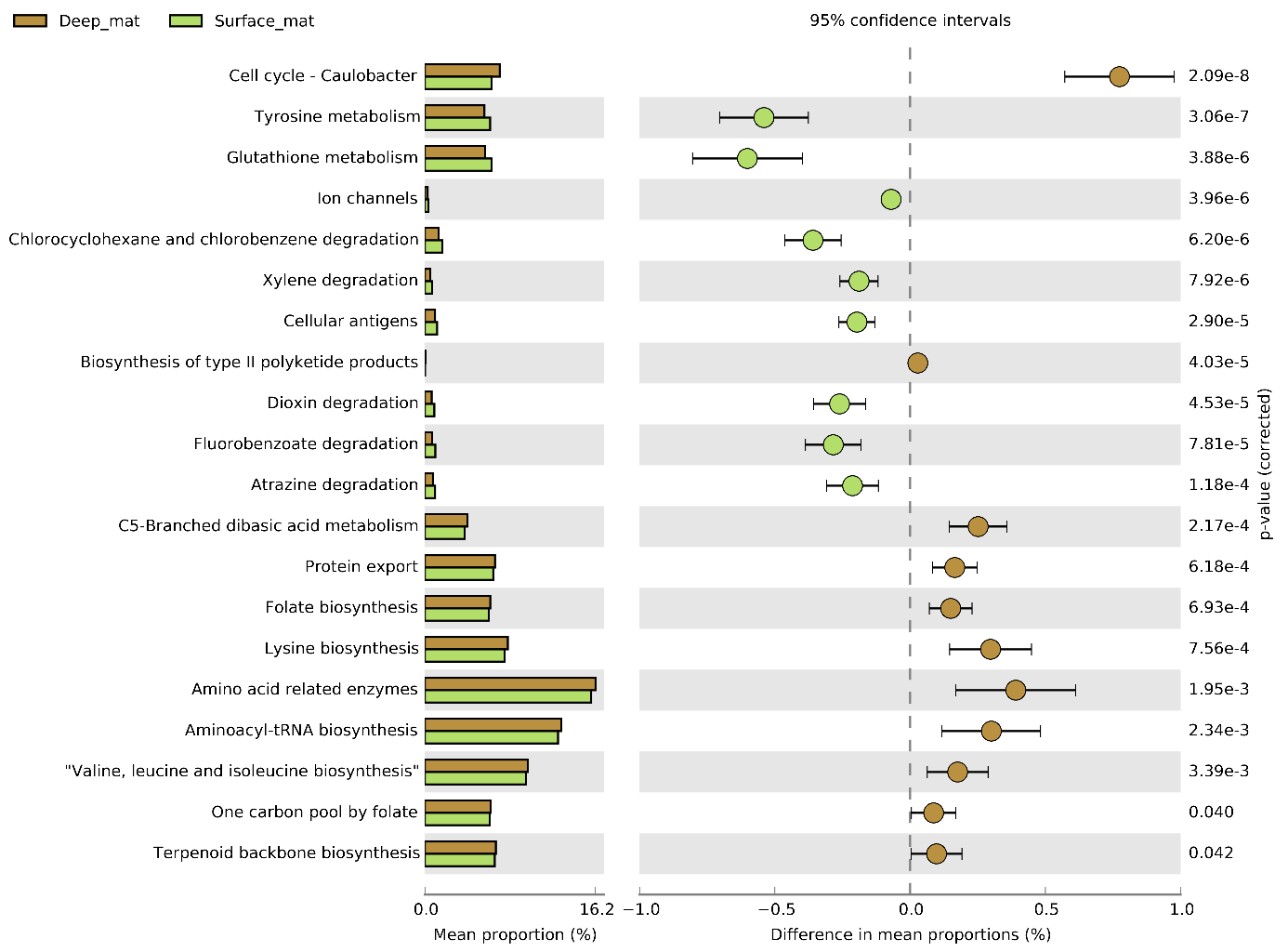

Supplementary Figure S7. Complete list of differences in the metabolic profiles comparing inferences for bacterial communities in microbial mats above (surface) and below (deep) perennial ice cover in Lake Vanda. Only statistically significant differences are displayed (P < 0.05).
